# Cryo-EM reveals the structural basis of subtype-specific, noncompetitive inhibition of the human P2X3 receptor

**DOI:** 10.64898/2026.01.03.697462

**Authors:** Zhixuan Zhao, Dong-Ping Wang, Xin Zhang, Yuan Gao, Hexin Xu, Xinyu Teng, Cheng Shen, Jirui Chen, Jinru Zhang, Ye Yu, Chang-Run Guo, Motoyuki Hattori

**Author notes:** These authors contributed equally: Zhixuan Zhao, Dong-Ping Wang.

## Abstract

P2X receptors are ATP-gated cation channels, and the P2X3 subtype plays crucial roles in peripheral sensory neurons, including in chronic pain and chronic cough. Accordingly, P2X3 has attracted substantial interest as a therapeutic target. Gefapixant, a negative allosteric modulator (NAM) of P2X3, has been approved in some countries for the treatment of chronic cough; however, its limited selectivity for P2X3 homomers over P2X2/P2X3 heteromers is associated with taste disturbance as a prominent adverse effect. These limitations have motivated the development of next-generation NAMs with improved subtype selectivity, but their subtype-specific allosteric inhibition mechanisms are unclear.

Here, we report the cryo-EM structure of the human P2X3 receptor in complex with ATP and the P2X3-selective next-generation NAM sivopixant, an investigational drug. Sivopixant binds to an allosteric site at the portal of the central pocket in the extracellular domain, and structure-based mutational analysis by electrophysiology identifies key residues required for sivopixant-dependent inhibition of human P2X3. Comparisons with P2X structures from other subtypes, together with gain-of-function mutants, define a structural basis for subtype-selective allosteric inhibition of the P2X3 receptor. Furthermore, structural comparisons with apo and ATP-bound open states of P2X3 receptors, together with molecular dynamics simulations, revealed that sivopixant expands the upper body domain to suppress the lower-body movements required for channel activation, thereby preventing channel opening even in the presence of ATP.

## Introduction

ATP is best known as the intracellular energy currency but also functions as an extracellular signaling molecule in many tissues via its release into the extracellular space (1, 2). P2X receptors are a family of ATP-gated cation channels that assemble as trimers from seven subtypes (3–7). ATP binding to P2X receptors induces conformational changes that open an ion-conducting pore, while the large extracellular domain also provides multiple pockets that can be targeted by endogenous regulators and small-molecule modulators (3, 4, 8). Given their widespread expression, P2X receptors contribute to diverse physiological processes, including synaptic transmission, nociception, inflammatory responses, and smooth muscle contractility (7, 8).

Among P2X subtypes, P2X3 receptors are expressed in peripheral sensory neurons (9–12). Consistent with this expression profile, P2X3 receptors have been associated with pathological roles in chronic pain and the cough reflex, motivating substantial interest in P2X3 as a therapeutic target (13, 14).

Clinical proof-of-concept for targeting P2X3 in chronic cough was established with AF-219 (gefapixant), a first-generation P2X3 negative allosteric modulator (NAM) that reduced cough frequency in refractory chronic cough (15). Gefapixant has since received marketing approval for chronic cough in Europe and Japan (16, 17). However, because gefapixant also inhibits P2X2/3 heteromers, adverse taste-related events have emerged as a clinically important limitation (18, 19); accordingly, gefapixant has not been approved by the U.S. FDA (16, 17). This connection is biologically plausible because ATP is a principal transmitter from taste buds to gustatory afferents, acting through P2X2/3 on sensory nerve fibers (20).

These limitations have driven the development of next-generation P2X3 modulators with improved selectivity over P2X2/3 heteromers (21, 22). S-600918 (sivopixant), a dioxotriazine-derived clinical candidate, was optimized for high potency at P2X3 with markedly higher selectivity over P2X2/3 (23). In a randomized clinical study in refractory chronic cough, sivopixant reduced cough frequency with a low incidence of taste disturbance, supporting the rationale that enhanced subtype selectivity may improve tolerability (24). In addition to sivopixant, other next-generation P2X3 modulators such as camlipixant and eliapixant have been developed with improved selectivity over P2X2/3 heteromers and have entered clinical trials (21, 25, 26).

Technological advances in structural biology have provided extensive structural information on P2X receptors and have deepened our understanding of their ATP-dependent gating mechanisms (27–36) , including those of P2X3 receptors. These studies have also highlighted that the orthosteric ATP-binding pocket is widely conserved across P2X subtypes, making it a poor target for subtype-selective modulation, and thereby further emphasizing the importance of NAMs for P2X receptors (37). For therapeutically targeted P2X receptors, particularly P2X4 and P2X7, a wide range of NAM-bound structures have been reported (38–43), revealing the structural basis of their subtype selectivity. In contrast, our understanding of the action of NAMs on P2X3 receptors (43–45), especially the mechanisms underlying next-generation modulators, remains limited. Whereas intensive structural studies and structure-based mutational analyses have been conducted for the first-generation NAM gefapixant (44, 46), for sivopixant, although *in silico* simulation and corresponding mutational analyses have been reported (47), a direct experimental structure capturing sivopixant bound to human P2X3 is lacking. Although a dog P2X3-camlipixant complex structure was recently reported, structure-based mutational analysis was not performed (45), obscuring the key structural determinants of subtype specificity. Overall, despite their clinical importance, the structural basis for the subtype selectivity of next-generation P2X3 receptor NAMs remains to be elucidated.

Likewise, structural information for ternary P2X-NAM-ATP complexes also remains limited, and to date only a ternary complex structure, panda P2X7 (pdP2X7) bound to its NAM A804598 and ATP, has been reported by X-ray crystallography (39). However, in the structural analysis of the panda P2X7-A804598-ATP complex, the entire cytoplasmic domain, which is required for full channel activation, was truncated (39), and the ternary structure was obtained by the crystal soaking method (39). These experimental conditions could influence the conformations captured in the structure; therefore, how NAMs prevent P2X receptor activation despite ATP binding is not yet fully understood.

To address these gaps, we report the cryo-EM structure of the human P2X3 receptor in complex with ATP and sivopixant. By combining the cryo-EM structure with electrophysiology and molecular dynamics simulations, we provide structural insights into the subtype specificity and noncompetitive inhibitory mechanism of the P2X3 receptor.

## Results

### Structure determination and overall structures

We purified the human P2X3 (hP2X3) protein, incubated it with sivopixant, and employed single-particle cryogenic electron microscopy (cryo-EM) to determine the structure of hP2X3 in complex with sivopixant. Cryo-EM data processing, particularly 3D variability analysis, yielded two distinct classes of EM maps (**Figs. 1 and 2 and Figs. S1 and S2**). The overall structures from the two cryo-EM maps are largely consistent and display a chalice-like trimeric assembly, in which each subunit comprises a large extracellular domain and two transmembrane (TM) helices (**Fig. 1**). This architecture matches the canonical P2X receptor fold and shows the characteristic dolphin-like shape (**Fig. 1C**), observed in previously reported P2X structures (29).

**Figure 1.**
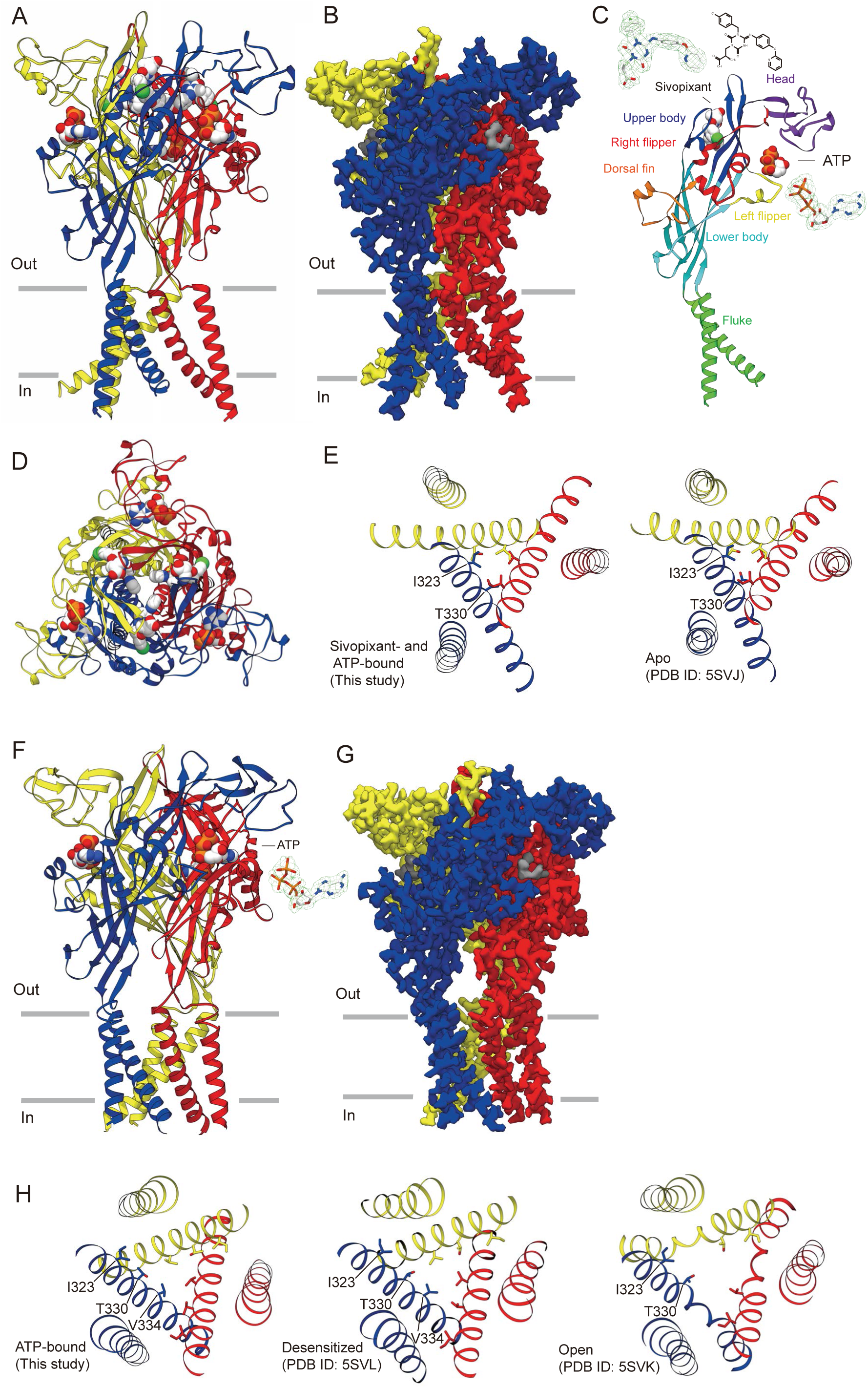
Overall structures. (A, B) Overall structure (A) and cryo-EM density map (B) of the sivopixant- and ATP-bound human P2X3 receptor viewed parallel to the membrane. Each subunit is colored distinctly. In A, the sivopixant and ATP molecules are shown in sphere representation. In B, the ligand densities are shown in gray. (C) The dolphin-shaped P2X3 subunit is colored differently according to each structural feature. The cryo-EM densities for sivopixant and ATP are shown. (D) Structure of the sivopixant- and ATP-bound human P2X3 receptor viewed perpendicular to the membrane from the extracellular side. (E) The transmembrane domain structures of the sivopixant- and ATP-bound P2X3 structure (left panel, this study) and the apo P2X3 structure (right panel, PDB ID: 5SVJ) viewed from the extracellular side. (F, G) The overall structure (F) and cryo-EM density map (G) of the ATP-bound human P2X3 receptor viewed parallel to the membrane. Each subunit is colored distinctly. In F, ATP molecules are shown in sphere representation, and the cryo-EM density for ATP is also shown. In G, ATP densities are shown in gray. (H) The transmembrane domain structures of the ATP-bound P2X3 structure (this study, left panel), the desensitized P2X3 structure (middle panel, PDB ID: 5SVL), and the open P2X3 structure (right panel, PDB ID: 5SVK) viewed from the extracellular side.

**Figure 2.**
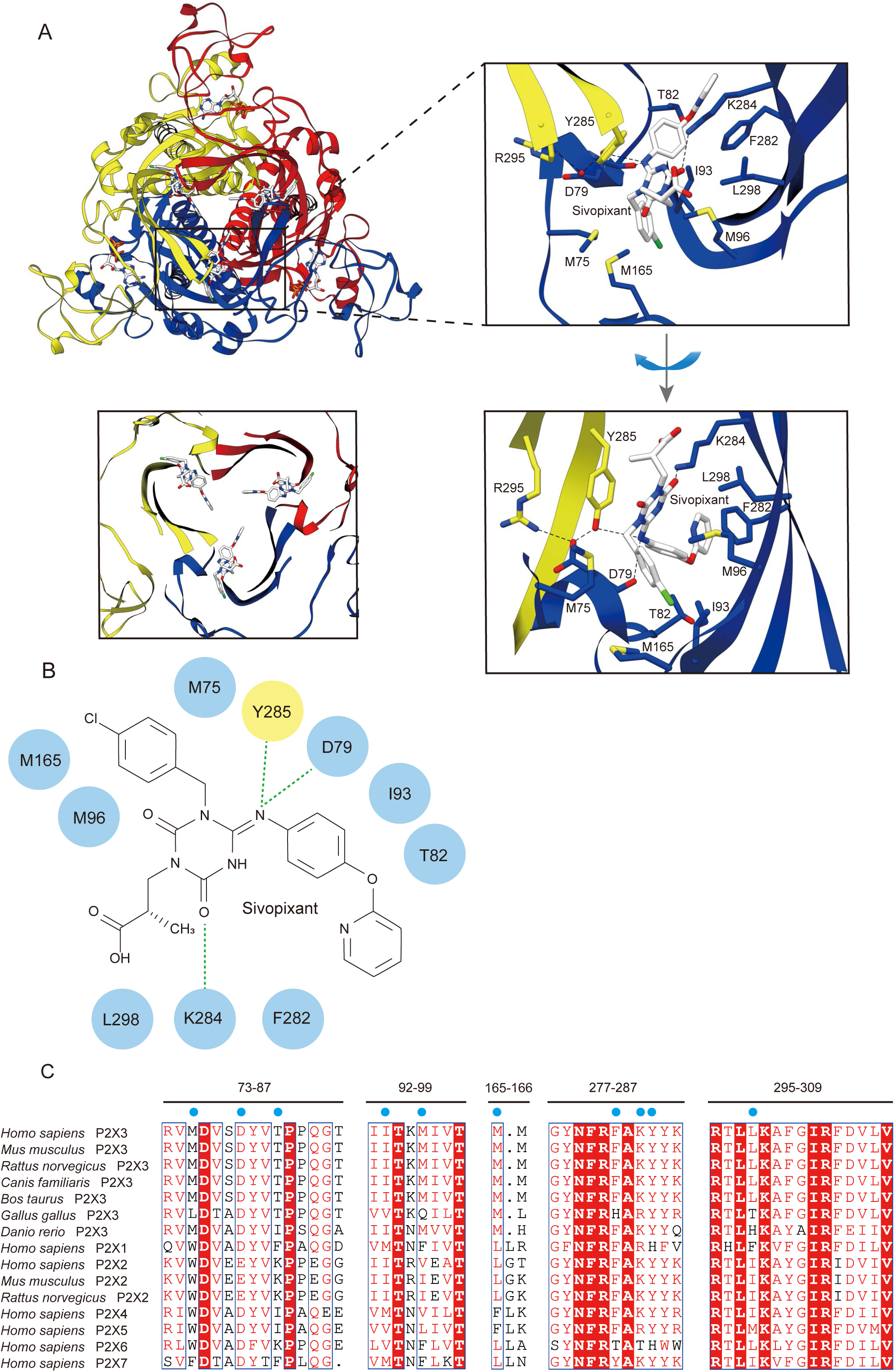
Sivopixant binding site. (A) Overall structure of the sivopixant- and ATP-bound human P2X3 receptor (upper-left panel) and a close-up view of the sivopixant binding site (lower-left panel) viewed perpendicular to the membrane from the extracellular side. Close-up views of the sivopixant binding site are shown from two different angles (upper-right and lower-right panels). The ligand molecules and amino acid residues involved in sivopixant binding are shown in stick representation. Dotted lines represent hydrogen bonds. (B) Schematic diagram of the interactions between P2X3 and sivopixant. Dotted lines represent hydrogen bonds. (C) Amino acid sequence alignment of P2X3 receptors *from Mus musculus* (Q3UR32.1), *Rattus norvegicus* (P49654.1), *Canis familiaris* (XP_038280235.1), *Bos taurus* (XP_059731161.1), *Gallus gallus* (NP_001384137.1), and *Danio rerio* (NP_571698.3); P2X receptors from *Homo sapiens* (P2X1: P51575.1; P2X2: Q9UBL9.1; P2X3: P56373.2; P2X4: Q99571.2; P2X5: Q93086.4; P2X6: O15547.2; and P2X7: Q99572.4) as well as the *Mus musculus* P2X2 receptor (Q8K3P1.2) and *Rattus norvegicus* P2X2 receptor (CAA71046.1). The residues involved in sivopixant binding are shown. The blue circles indicate the residues shown in B.

However, as an important difference, in one map at 3.34 Å resolution, we observed residual cryo-EM density consistent with that of sivopixant and ATP (**Fig. C**), whereas in the other map at 2.95 Å resolution, we observed residual density consistent with that of ATP alone (**Fig. 1F**). Owing to the high affinity of P2X3 for ATP, ATP observed in these structures likely originated from endogenous ATP in cells that remained bound during purification (31, 45).

Consistent with the cobinding of ATP and sivopixant to P2X3 in our cryo-EM structure, whole-cell patch-clamp recordings of hP2X3 showed that the sivopixant-dependent inhibition of ATP-dependent channel currents was independent of the ATP concentration (**Figs. 3A and 3B**), indicating that sivopixant binding to P2X3 does not compete with ATP. Moreover, in our sivopixant- and ATP-bound structure, the TM domain adopted a previously-reported apo, closed conformation, with Ile323 and Thr330 forming a pore constriction (31) (**Fig. 1E**). In contrast, in our ATP-bound structure, the TM domain adopted a conformation resembling the ATP-bound desensitized state, with Val334 forming the pore constriction (31) (**Fig. 1H**). These observations suggest that sivopixant inhibits the channel not by stabilizing the desensitized state, but rather by stabilizing a closed state, even in the presence of ATP.

**Figure 3.**
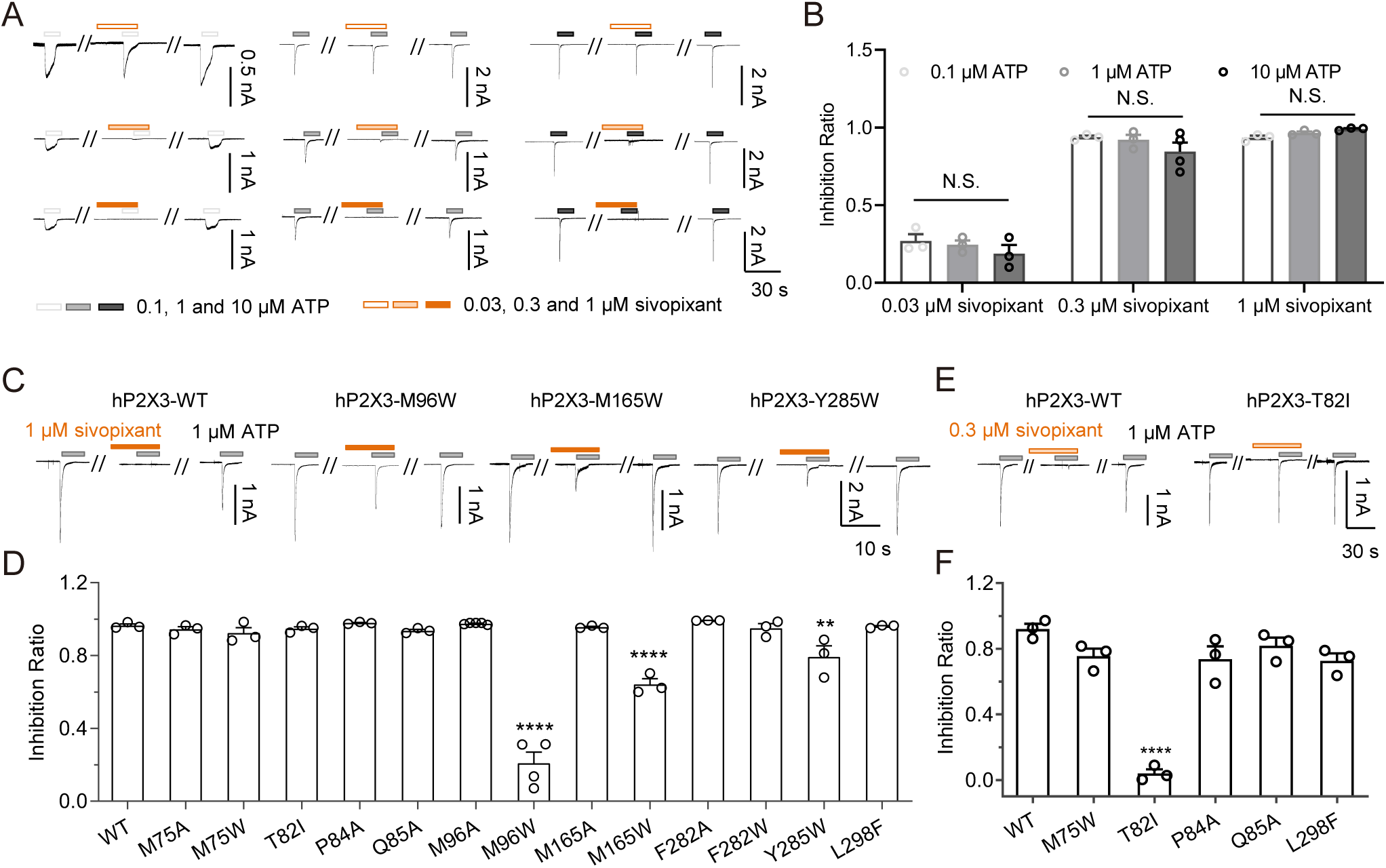
Mutational analysis. (A) Representative current traces of the effects of sivopixant on human P2X3 currents at different ATP concentrations. (B) Effects of sivopixant on ATP (0.1, 1 and 10 µM)-evoked currents of human P2X3 (mean ± SEM, n = 3-4). (C, E) Representative current traces of sivopixant effects at 1 μM (C) and 0.3 μM (E) on ATP-evoked currents of human P2X3 and its mutants (C: M96W, M165W, and Y285W; E: T82I). (D, F) Effects of 1 μM (D) and 0.3 μM (F) sivopixant on the ATP-evoked currents of human P2X3 and its mutants (mean ± SEM, n = 3-5). Two-way ANOVA followed by Tukey‘s multiple comparisons test (B) and one-side one-way ANOVA followed by post hoc test (D, F), **p < 0.01, ****p < 0.0001 vs. WT). The hP2X3 WT data shown in Figs. 3 and 5 were obtained from the same cells and are shown in the respective panels for comparison.

### Sivopixant binding site

In the sivopixant- and ATP-bound structure, sivopixant binds at each trimeric interface within the extracellular domain (**Figs. 1A and 1D**). In the dolphin representation, sivopixant is located in the upper body region and is therefore distant from the ATP-binding site (**Fig. 1C**). Notably, a corresponding cavity space is present not only in P2X3 but also in other P2X subtypes including P2X4 and P2X7 (38–42), where the cavity space serves as a hotspot for small-molecule modulation and has been referred to as the portal of the central pocket (PCP) (48).

The carboxyl group of sivopixant faces the outer side of the PCP, whereas the pyridine moiety faces the inner side (**Fig. 2A**). Sivopixiant binding is mediated mainly by hydrophobic contacts involving Met75, Ile93, Met96, Met165, Phe282, and Leu298 from one subunit and Tyr285 from the neighboring subunit (**Figs. 2A and 2B**). In addition, the main-chain carbonyl of Asp79 and the side chain of Tyr285 form hydrogen bonds with a nitrogen atom of the triazine ring, and the side chain of Lys284 forms a hydrogen bond with the carbonyl oxygen of the triazine ring (**Figs. 2A and 2B**). Furthermore, intersubunit salt bridges between Asp79 of one subunit and Arg295 of the neighboring subunit, as well as hydrogen bonds between Asp79 in one subunit and Tyr285 of the neighboring subunit, appear to stabilize the pocket architecture (**Fig. 2A**).

Among the residues implicated in sivopixant binding, Asp79, Thr82, Met96, Lys284, and Arg295 have been suggested to be important for sensitivity to the sivopixant analog 3-(4-([3-chloro-4-isopropoxyphenyl]amino)-3-(4-methylbenzyl)-2,6-dioxo-3,6-dihydro-1,3,5-triazin-1(2H)-yl)propanoic acid (DDTPA) and/or to sivopixant itself (47). However, for many of the additional residues that contact sivopixant in our structure, mutational validation has not yet been performed. This is likely because our previous *in silico* modeling placed DDTPA toward the outer periphery of this allosteric pocket (**Fig. S3A**), and the subsequent mutagenesis was designed on the basis of the predicted binding pose.

### Structure-based mutational analysis

We performed structure-based mutational analyses based on our experimental structure (M75A, M75W, T82I, P84A, Q85A, M96A, M96W, M165A, M165W, F282A, F282W, Y285W, and L298F) (**Fig. 3**). We introduced alanine substitutions to attenuate side chain-mediated interactions with sivopixant. We also introduced bulkier side chain substitutions (to Trp or Phe) to create steric interference with sivopixant and thereby weaken its binding. In addition, T82I was designed to introduce a residue type commonly found at the corresponding position in other P2X subtypes (**Fig. 2C**).

First, we tested all the mutants for the inhibition of ATP-dependent channel activation by 1 μM sivopixant (**Figs. 3C and 3D**). Compared with the wild type, M96W, M165W, and Y285W showed reduced sensitivity to sivopixant. Next, for a subset of mutants that did not show an obvious change in sensitivity to 1 μM sivopixant (M75W, T82I, P84A, Q85A, and L298F), we performed patch-clamp recordings at a lower concentration of 0.3 μM sivopixant. Under this condition, T82I exhibited a significant reduction in sensitivity (**Figs. 3E and 3F**). Together, these structure-based mutational analyses revealed residues that contribute to sivopixant sensitivity.

### Subtype specificity

To gain structural insights into the high subtype selectivity of sivopixant, we superposed our sivopixant-bound hP2X3 structure with previously-reported structures of other P2X subtypes (**Fig. 4**). Integrating the structural comparisons with our mutational analysis results and a sequence alignment across P2X subtypes, we focused on five residues: Asp79, Thr82, Met96, Met165, and Tyr285 for further analysis (**Figs. 2C, 3 and 4**). Among these, Asp79 has been implicated in prior mutational studies (47), whereas Thr82, Met96, Met165, and Tyr285 were identified as the residues involved in sivopixant sensitivity in our present analyses (**Fig. 3**).

**Figure 4.**
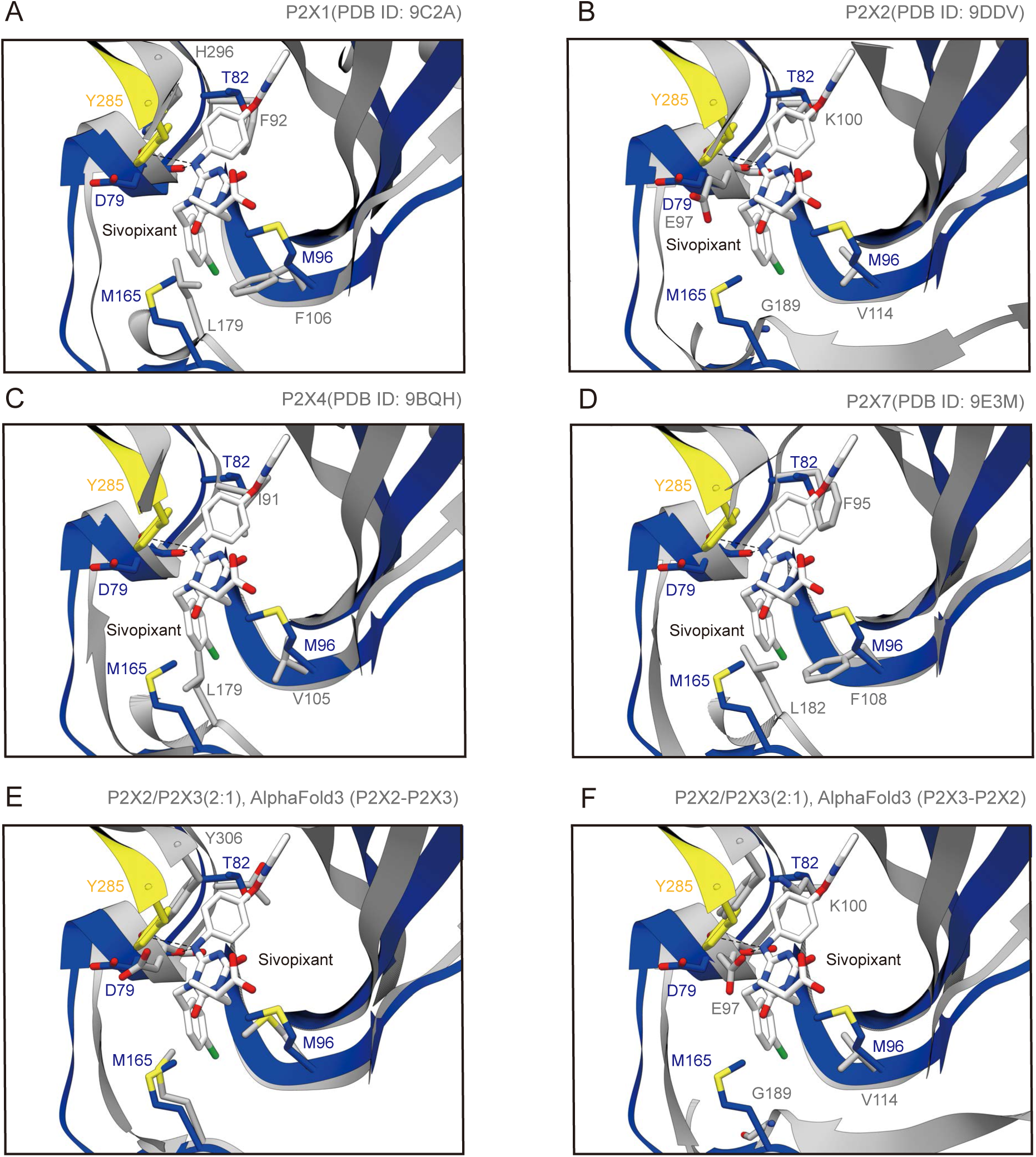
Subtype specificity. (A-F) Close-up views of the sivopixant binding site of the sivopixant- and ATP-bound P2X3 structure (in this study, yellow and blue). The human P2X1 structure (PDB ID: 9C2A) (A), the human P2X2 structure (PDB ID: 9DDV) (B), the human P2X4 structure (PDB ID: 9BQH) (C), the human P2X7 structure (PDB ID: 9E3M) (D), and the predicted heterotrimer structure formed by two P2X2 subunits and one P2X3 subunit (AlphaFold3, ipTM=0.71) (E, F) are superposed onto the P2X3 structure and shown in gray. In E, the gray chain superposed onto the yellow chain is the P2X2 subunit, while the gray chain superposed onto the blue chain is the P2X3 subunit. In F, the gray chain superposed onto the yellow chain is the P2X3 subunit, while the gray chain superposed onto the blue chain is the P2X2 subunit.

First, comparison of the sivopixant-bound hP2X3 structure with the previously reported human P2X1 structure revealed that Thr82, Met96, Met165, and Tyr285 in hP2X3 are replaced by Phe92, Phe106, Leu179, and His296 in human P2X1, respectively (**Fig. 4A**). Similarly, residues corresponding to Asp79, Thr82, Met96, and Met165 in hP2X3 are substituted in P2X2 by Glu97, Lys100, Val114, and Gly189, respectively (**Fig. 4B**). In P2X4 (**Fig. 4C**), the residues corresponding to Thr82, Met96, and Met165 in hP2X3 are Ile91, Val105, and Leu179, respectively, and in P2X7, those residues are Phe95, Phe108, and Leu182, respectively (**Fig. 4D**).

No experimental structures have been reported for P2X2/P2X3 heterotrimers; therefore, we generated predicted structural models of human P2X2/P2X3 heteromers with different stoichiometries (P2X2/P2X3 (2:1) and P2X2/P2X3 (1:2)) using AlphaFold3 (49). Both models exhibited similar subunit interfaces (**Figs. 4E, 4F and S4**); therefore, we used the P2X2/P2X3 (2:1) model for subsequent discussion. Among the two types of subunit interfaces, one interface (P2X2-P2X3) did not differ in the corresponding region (**Fig. 4E**), whereas the other interface (P2X3-P2X2) displayed clear differences derived from P2X2, with the residues corresponding to Asp79, Thr82, Met96, and Met165 in hP2X3 replaced by Glu97, Lys100, Val114, and Gly189 (**Fig 4F**). Collectively, these analyses indicate that these five residues (Asp79, Thr82, Met96, Met165, and Tyr285) constitute a hotspot of intersubtype sequence divergence.

### Gain of function mutants

Finally, on the basis of the abovementioned insights, we designed gain-of-function (GOF) mutants of human P2X1 and P2X2 to confer sivopixant sensitivity for understanding subtype specificity (**Fig. 5**). Our previously designed P2X2 GOF mutant (E97D/K100T/E103Q/G105T) did not confer sivopixant sensitivity to the P2X2 homotrimer but could confer partial sensitivity to P2X2/P2X3 heteromers (47). In this previous mutant, the targeted positions corresponding to E103Q (Gln85 in hP2X3) and G105T (Thr87 in hP2X3) do not directly contact sivopixant in our structure but are located near the binding pocket and could influence the shape of the pocket. In contrast, E97D (Asp79 in hP2X3) and K100T (Thr82 in hP2X3) correspond to two of the five residues we mentioned above (Asp79, Thr82, Met96, Met165, and Tyr285) and are positioned to directly affect sivopixant binding. The remaining three residues (Met96, Met165, and Tyr285) were not incorporated into earlier gain-of-function designs, because an experimental structure of the binding mode was not available.

**Figure 5.**
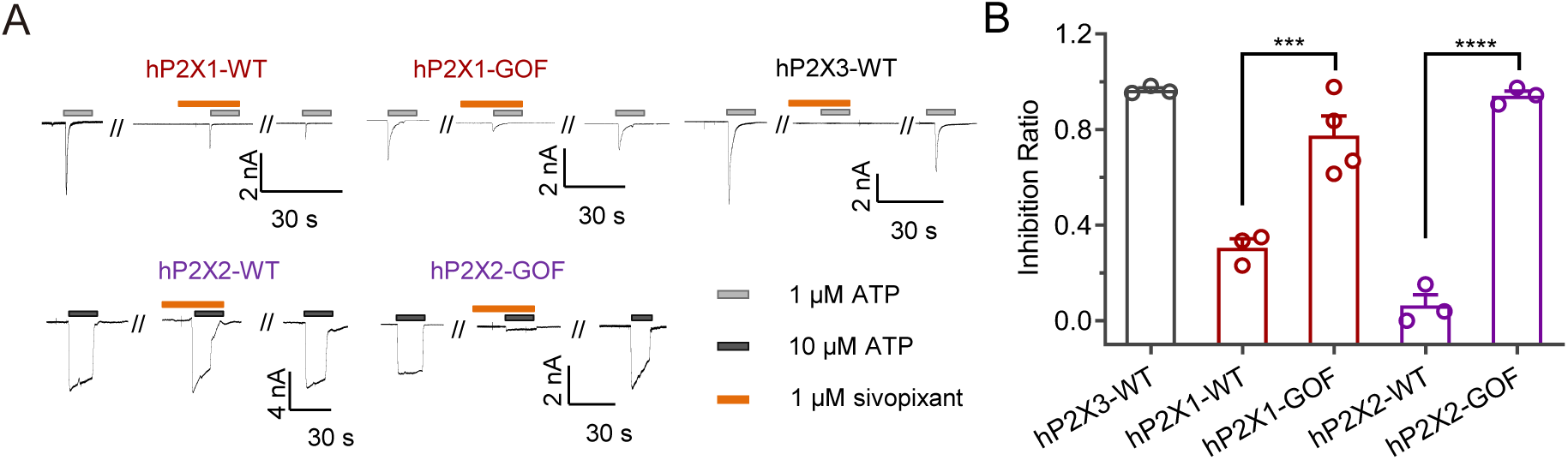
Gain of function mutants. (A) Representative current traces of the effects of sivopixant on ATP-evoked currents of human P2X3 wild type (WT) and its gain-of-function mutant (GOF). (B) Effects of sivopixant on ATP-evoked currents of human P2X3 and its mutant (mean ± SEM, n = 3-4). (One-way ANOVA followed by Tukey‘s multiple comparisons test, ***p < 0.001 hP2X1GOF vs. hP2X1-WT, ****p < 0.0001 hP2X2GOF vs. hP2X2-WT). The hP2X3 WT data shown in Figs. 3 and 5 were obtained from the same cells and are shown in the respective panels for comparison.

Accordingly, starting from the P2X2 E97D/K100T/E103Q/G105T background, we introduced additional substitutions to match the five key residues to their hP2X3 counterparts in human P2X1 and P2X2 (hP2X1 GOF: F92T/D97T/F106M/L179M/H296Y; hP2X2 GOF: E97D/K100T/E103Q/G105T/V114M/G189M). Strikingly, whereas the wild-type hP2X2 homomer was insensitive to sivopixant, the new GOF mutant clearly displayed sivopixant sensitivity (**Fig. 5**). In addition, although wild-type hP2X1 exhibited only weak sensitivity to 1 μM sivopixant, the sensitivity of the GOF mutant significantly increased (**Fig. 5**). Together, these results show that the five residues implicated in our comparisons (Asp79, Thr82, Met96, Met165, and Tyr285) play a critical role in the subtype selectivity of sivopixant.

### Structural comparison

To address how sivopixant prevents P2X3 activation despite ATP binding, we compared our hP2X3 structure in complex with sivopixant and ATP with previously reported P2X3 structures in the apo, closed state and ATP-bound, open state (**Fig. 6**). First, the ATP-binding mode in our structure is essentially the same as that in the ATP-bound, open state (**Figs. 6B and 6C**), which is consistent with the noncompetitive action of sivopixant on P2X3 (**Figs. 3A and 3B**). Consistent with ATP binding, we observed ATP-dependent upward motion of the dorsal fin and downward motion of the left flipper to some extent (**Fig. 6A**), even in the presence of sivopixant. The upward motion of the dorsal fin and downward motion of the left flipper are known to be important for channel activation (50), since these motions are normally coupled with the expansion of the lower body domain, leading to channel opening (**Figs. 6A**) (28). However, the conformational changes in the lower body domain, particularly those near the TM domain, were largely absent in the sivopixant-bound structure (**Fig. 6A**). Accordingly, the ion-conducting pore remains closed (**Fig. 1E**). The lack of conformational changes in the lower body domain is likely due to sivopixant-dependent movement of the upper body domain (**Fig. 6A**). The upper body domain serves as a pivot for lower-body movements (28); thus, expansion of the upper body may prevent it from functioning as an effective pivot, instead exerting force in the opposite direction and, thereby hindering lower-body expansion (**Figs. 6D and 6E**). This notion is consistent with the previous mutational cross-linking analysis of the upper body domain in P2X7 (51).

**Figure 6.**
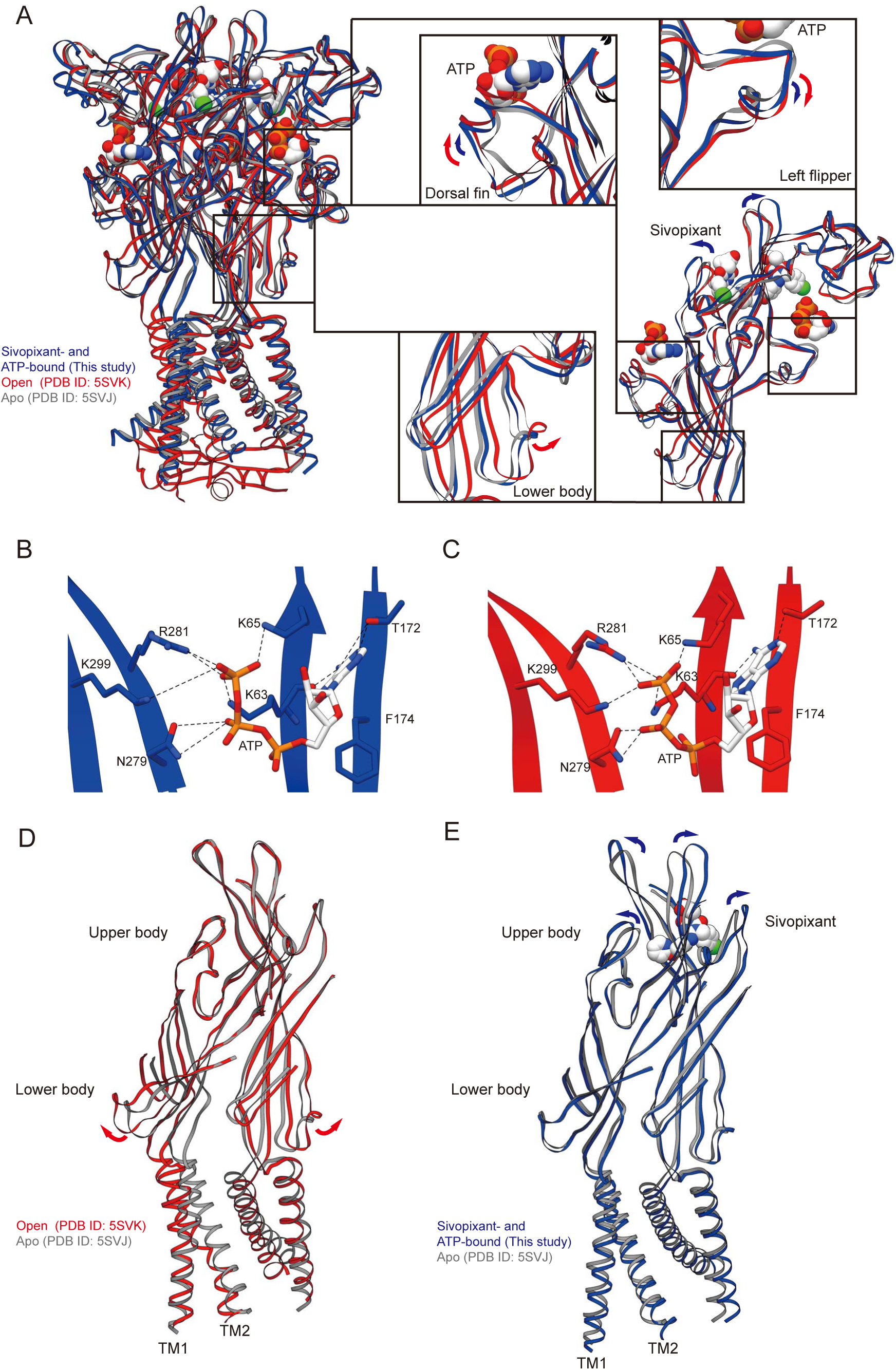
Structural comparison. (A) Superimposition of the sivopixant- and ATP-bound P2X3 structure (in this study, blue) and the ATP-bound, open P2X3 structure (PDB ID: 5SVK, red) onto the apo, closed P2X3 structure (PDB ID: 5SVJ, gray) viewed parallel to the membrane. Close-up views of the dorsal fin, left flipper, and lower body domains are also shown. The arrows indicate conformational changes in the open P2X3 structure (red) and the sivopixant- and ATP-bound P2X3 structure (blue). (B, C) Close-up view of the ATP binding site of the sivopixant- and ATP-bound P2X3 structure (in this study, blue) (B) and the open P2X3 structure (PDB ID: 5SVK, red) (C). The ATP molecules and amino acid residues involved in ATP binding are shown in stick representation. Dotted lines represent hydrogen bonds. (D, E) Superimposition of the open P2X3 structure (PDB ID: 5SVK, red) (D) and the sivopixant- and ATP-bound P2X3 structure (this study, blue) (E) onto the apo P2X3 structure (PDB ID: 5SVJ, gray). Only the transmembrane and body domains from the two subunits in the foreground are shown.

To test whether the structural changes in the upper body domain observed in our cryo-EM structure are dependent on sivopixant binding, we performed molecular dynamics (MD) simulations using the cryo-EM structure in complex with sivopixant and ATP as starting models, as well as models in which sivopixant or ATP was deleted, and a model in which both were deleted (**Fig. 7**). In all MD runs, the overall structures and ligand binding were largely stable **(Fig. S5**). Consistent with our model, the expansion of the upper body domain were largely stable either with or without ATP throughout the MD simulations of the sivopixant-bound structures (**Figs. 7B and 7C**), whereas the removal of sivopixant led to shorter distances between the Cα atoms of Glu289 in the upper body domain from two adjacent subunits of the trimer, indicating a closing motion of the upper body domain in the absence of sivopixant (**Figs. 7D and 7E**).

**Figure 7.**
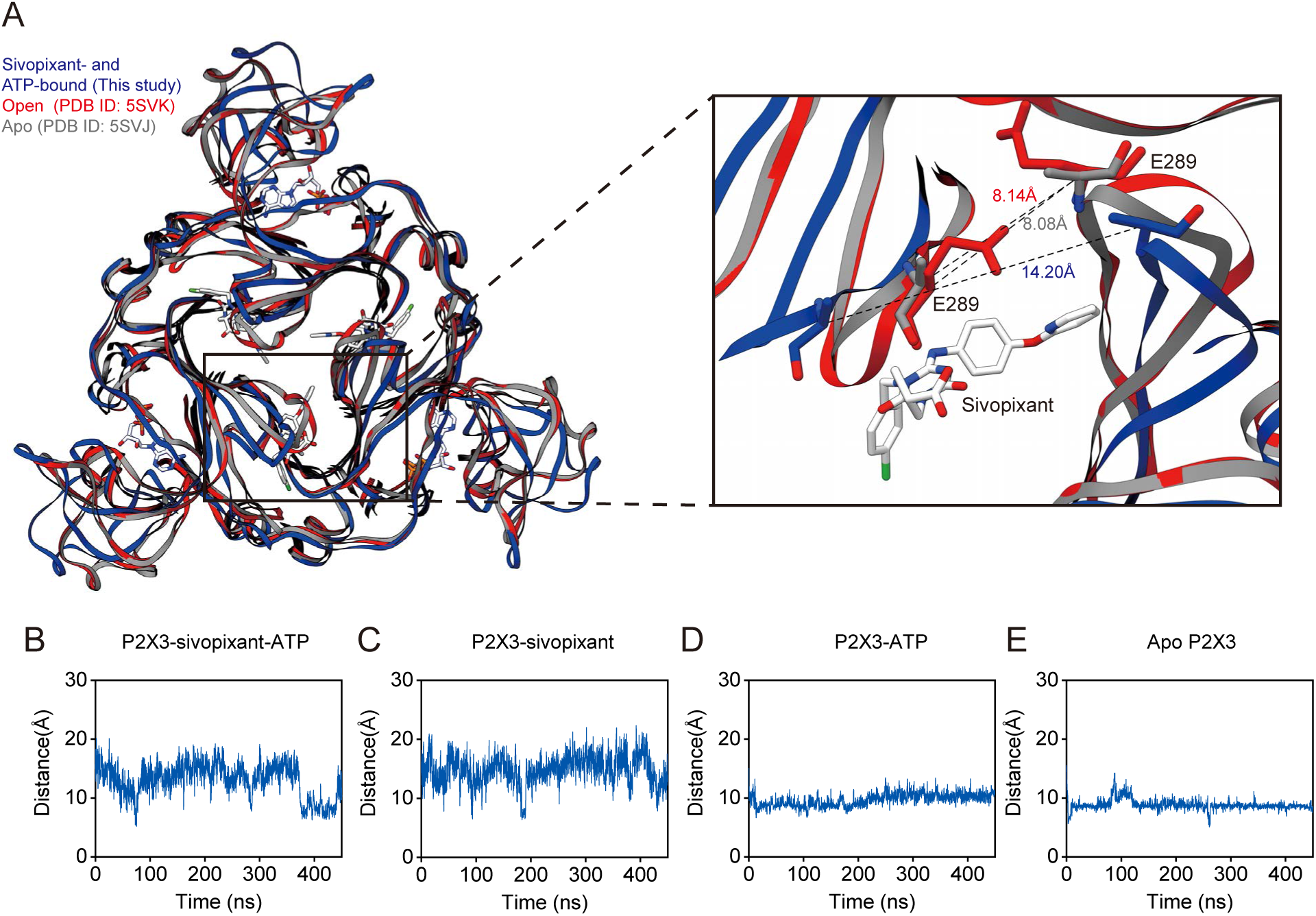
MD simulations. (A) Close-up view of the sivopixant binding site of the P2X3 receptor. Superimposition of the sivopixant- and ATP-bound P2X3 structure (this study, blue) and the open P2X3 structure (PDB ID: 5SVK, red) onto the apo P2X3 structure (PDB ID: 5SVJ, gray) viewed perpendicular to the membrane from the extracellular side. Dotted lines indicate the distance (Å) between the Cα atoms of Glu289 in two adjacent subunits. (B-E) MD simulations using the sivopixant- and ATP-bound structure with both retained (B), ATP deleted (C), sivopixant deleted (D) and both deleted (E) as starting models. The distance plots of Cα atoms between Glu289 of two adjacent subunits are shown. The average distances in the trimer are shown.

Taken together, these results suggest that sivopixant binding induces structural changes in the upper body domain, which in turn leads to uncoupling between the upper and lower body domains and thereby prevents channel activation even with the binding of ATP (**Fig. 8**).

**Figure 8.**
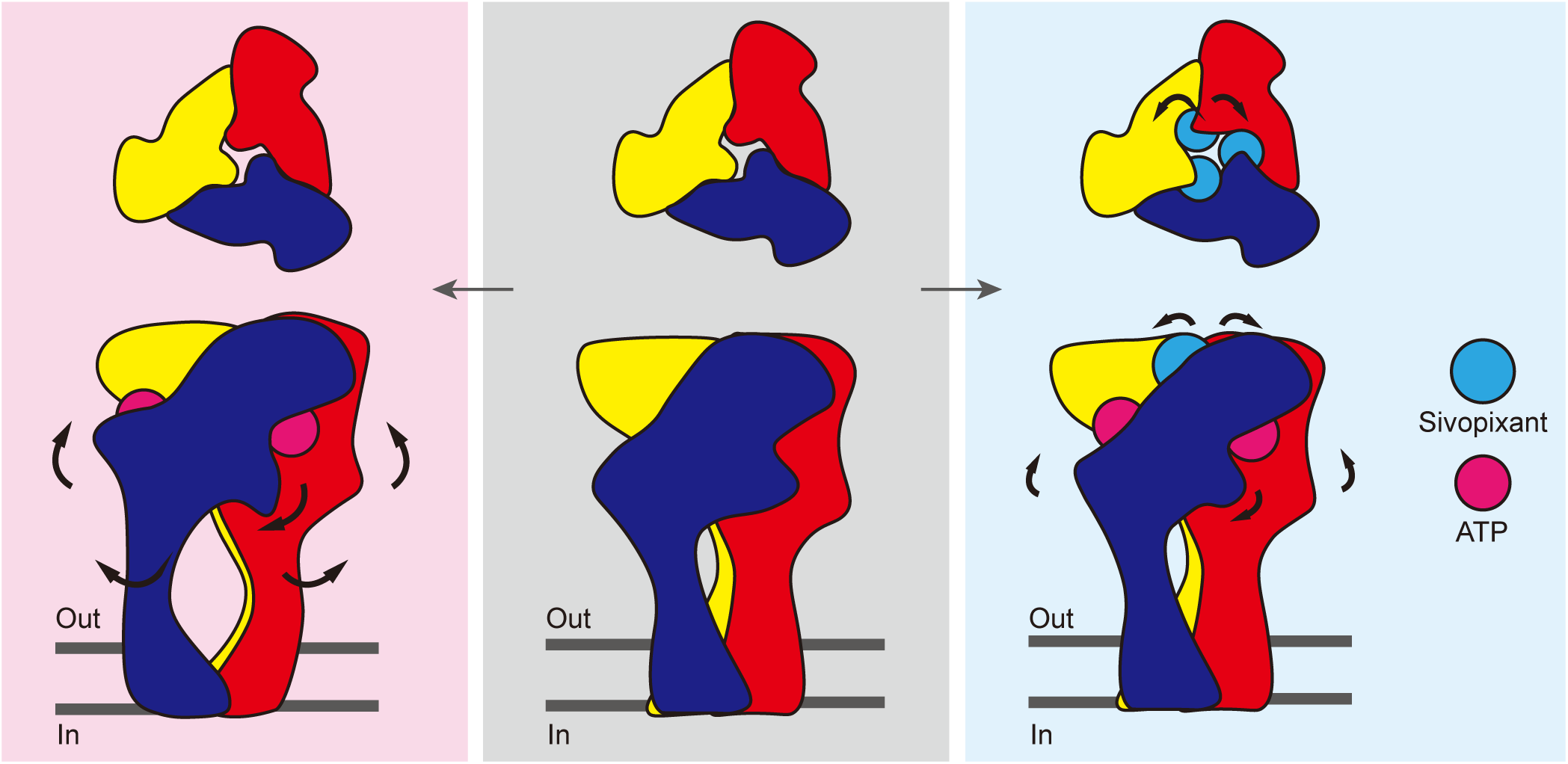
Proposed noncompetitive inhibition mechanism. Cartoon diagrams illustrating the conformational changes of P2X3 from the apo state (middle) to the ATP-bound, open state (left) and the sivopixant- and ATP-bound, closed state (right). The arrows indicate conformational changes between two states.

## Discussion

In this work, we determined the cryo-EM structure of the human P2X3 receptor in complex with ATP and sivopixant (**Figs. 1 and 2**) and performed structure-based mutational analysis of the ligand binding site using patch-clamp recordings (**Fig. 3**). The binding pose of sivopixant in our cryo-EM differs substantially from that of a sivopixant analog based on our previous *in silico* modeling (47)(**Fig. S3A**). This discrepancy likely reflects the difficulty of predicting ligand binding starting from an apo structure, in which the allosteric site is narrower than in the ligand-bound state (**Fig. S3A**) and therefore cannot accommodate the ligand deeply within the pocket. Consistently, when we attempted to predict human P2X3 in complex with sivopixant using Boltz-2 (52), an AlphaFold3-derived model capable of small-molecule complex prediction, sivopixant was not placed at the allosteric site, but at unrelated sites, such as the ATP-binding pocket or the TM domain. Even when we imposed restraints on the five residues important for the subtype specificity (Asp79, Thr82, Met96, Met165, and Tyr285) in the allosteric site, the predicted pose was oriented opposite to that observed in the cryo-EM structure (**Fig. S3B**). Together, these results show that despite recent advances in AI-based structure prediction, accurate prediction of small-molecule binding remains challenging (53).

Structural comparisons with other P2X subtypes, together with patch-clamp recordings of gain-of-function mutants, revealed key residues for subtype-specific inhibition (**Figs. 4 and 5**). This allosteric site corresponds to the portal of the central pocket (PCP), a hotspot for allosteric modulators across multiple P2X receptors (48). Notably, in previously reported P2X structures in complex with subtype-selective allosteric modulators (**Fig. S6**), many of the corresponding residues are involved in ligand binding, suggesting that sequence diversity at these positions provides a rational basis for designing subtype-selective allosteric modulators. Moreover, interspecies sequence differences within the PCP can cause species-dependent differences in compound sensitivity, particularly between humans and rodents, thereby complicating preclinical studies, as exemplified by the presence of Ile315 in human P2X4 and Val312 in human P2X7 (48, 54). In contrast, the residues involved in sivopixant binding are fully conserved among human, mouse, and rat P2X3 (**Fig. 2**). Therefore, our work provides a foundation for designing compounds that are subtype selective and not affected by interspecies differences.

Our cryo-EM structure captures a ternary complex containing sivopixant and ATP; therefore, our structure also provides insight into how a NAM can inactivate the channel despite ATP binding. Structural comparisons with apo and ATP-bound open states, together with molecular dynamics simulations, revealed sivopixant-dependent expansion of the upper body domain (**Figs. 6-8**). Although we observed ATP-dependent conformational changes in the dorsal fin and left flipper (**Fig. 6A**), which are normally associated with activation, the upper body domain, which does not directly participate in ATP binding, undergoes sivopixant-dependent expansion to suppress the expansion of the lower body domain and thereby inhibits channel opening. This inhibitory effect of lower body domain expansion on channel activation is consistent with previous mutational analysis (51). On the other hand, in the previously reported crystal structure of the panda P2X7-NAM-ATP ternary complex (39), such NAM-dependent upper-body expansion was not observed. One possible explanation is that the experimental conditions used for structure determination, such as truncation of the entire cytoplasmic domain, which is required for full channel activation, and the soaking method, could have influenced the conformations captured in the previous structure.

Taken together, our work provides a structural basis for the subtype specificity and noncompetitive inhibition of the human P2X3 receptor by the investigational drug sivopixant, which may facilitate the rational design and optimization of next-generation P2X3 negative allosteric modulators for chronic cough and other P2X3-related sensory disorders.

## Methods

### Protein expression and purification

The DNA sequence encoding the functional construct of human P2X3 for the structural studies (31) was synthesized by Genewiz Inc. (Suzhou, China) and subcloned into a pFastBac vector containing a Twin-Strep tag, an EGFP tag, and a Tobacco etch virus (TEV) protease cleavage site at the N-terminus. DH10Bac *Escherichia coli* cells were used as the host for bacmid recombination. Sf9 cells were cultured in suspension at 27 °C in SIM SF culture medium (Sino Biological, China) and routinely passaged every other day. The initial recombinant baculovirus was generated by transfecting adherent Sf9 cells with bacmid DNA using FuGENE HD reagent (Promega, USA) and was used to infect Sf9 cells for virus amplification. The EGFP-fused P2X3 protein was overexpressed in Sf9 suspension cultures at 27 °C for 60 hours post-infection. Cells were collected and lysed by ultrasonication in TBS buffer (50 mM Tris-HCl, pH 8.0, and 150 mM NaCl) supplemented with 1mM phenylmethylsulfonyl fluoride (PMSF), 5.2 μg/mL aprotinin, 2 μg/mL leupeptin, and 1.4 μg/mL pepstatin A. The supernatant was collected after centrifugation at 8,000 × g for 20 min and then ultracentrifuged at 185,000 × g for 1 hour. The membrane fraction pellets were solubilized in solubilization buffer [50 mM Tris-HCl, pH 8.0, 150 mM NaCl, 15% glycerol, and 2% n-dodecyl-beta-d-maltopyranoside (DDM) (Anatrace, USA)] supplemented with 1mM PMSF, 5.2 μg/mL aprotinin, 2 μg/mL leupeptin, 1.4 μg/mL pepstatin A, and 0.2 unit/mL apyrase (Sigma, USA), and stirred for 1.5 hour. The solubilized mixture was ultracentrifuged at 185,000 × g for 1 hour. The supernatant was loaded onto Streptactin Beads 4FF beads (Smart-Lifesciences, China) pre-equilibrated with wash buffer (50 mM Tris-HCl, pH 8.0, 150 mM NaCl, 5% glycerol, and 0.05% DDM) and stirred for 1 hour. The resin was subsequently washed with 12 CV of the wash buffer. The proteins were eluted with elution buffer (50 mM Tris-HCl, pH 8.0, 150 mM NaCl, 5% glycerol, 0.05% DDM, and 2.5 mM desthiobiotin) and cleaved with TEV protease to remove EGFP and the Twin-Strep tag by dialysis overnight in dialysis buffer (150 mM NaCl, 20 mM HEPES, pH 7.5, and 0.025% DDM). The cleaved protein was applied to a Superdex 200 Increase 5/150 GL column (Cytiva, USA) pre-equilibrated with gel filtration buffer [150 mM NaCl, 20 mM HEPES, pH 7.5, 0.001% lauryl maltose neopentyl glycol (Anatrace, USA) and 0.0001% cholesteryl hemisuccinate (CHS)]. Peak fractions were collected and concentrated to 4 mg/mL using an Amicon Ultra 50 kDa cutoff (Merck Millipore, USA). All purification processes were carried out at 4 °C.

### Cryo-EM data acquisition

For grid preparation, the P2X3 protein was mixed with 0.1 mM Sivopixant (MedChemExpress, China). Following ultracentrifugation at 185,000 × g for 20 min, the protein was applied to holey carbon-film grids (Quantifoil, Germany, Au R1.2/1.3 μm size/hole space, 300 mesh) that had been glow-discharged for 60 seconds, blotted using a Vitrobot (ThermoFisher Scientific, USA) at 100% humidity and 4 °C, and plunge-frozen in liquid ethane cooled by liquid nitrogen. Cryo-EM movies were acquired using a Falcon 4i camera (Thermo Fisher Scientific, USA) equipped on a Titan Krios (Thermo Fisher Scientific, USA) electron microscope at an acceleration voltage of 300 kV at a magnification of 130,000× for the dataset. A total of 7,464 movies were collected with a total dose of 40 electrons per Å^2^, a pixel size of 0.959 Å, and a defocus range of -1.2 to -1.8 μm.

### Cryo-EM data processing

All data processing steps were performed in CryoSPARC (55). Movies were motion-corrected and contrast transfer function parameters were estimated. Particles were extracted with a box size of 288 pixels, followed by two rounds of template-based particle picking and 2D classification. An ab initio reconstruction was then generated with C3 symmetry imposed. Well-defined subsets were selected and subjected to two rounds of heterogeneous refinement and nonuniform refinement (56). After heterogeneous refinement using two 3D volumes obtained from 3D variability analysis (57), the two classes were further processed by nonuniform refinement, two rounds of local refinement, map sharpening, and local resolution estimation. The final resolution for P2X3 with ATP was 2.95 Å from 71,855 particles, whereas the final resolution for P2X3 with both sivopixant and ATP was 3.34 Å from 46,550 particles (**Table S1**).

### Model building and refinement

All atomic models for the sivopixant- and ATP-bound structure and the ATP-bound structure were built in Coot (58) using previously reported P2X3 structures in the apo state (PDB ID: 5SVJ) and the ATP-bound, desensitized state (PDB ID: 5SVL), respectively. After manual model adjustment in Coot, the structures were refined by real-space refinement in PHENIX (59) (**Table S1**). Structure figures were generated using UCSF Chimera (60). The sequence-alignment figure was generated using Clustal Omega (61) and ESPript 3.0 (61). Predicted structural models of human P2X2/3 heteromers were generated using AlphaFold3 (49). Sivopixant binding prediction was performed using Boltz-2 (52).

### Electrophysiology

The hP2X3 plasmid was purchased from Open Biosystems (USA). cDNAs for hP2X1 and hP2X2 were synthesized by BGl Genomics (China) and subcloned into the pEGFP-N1 vector. All mutations were generated using the KOD-Plus-Mutagenesis Kit (TOYOBO, Japan) and verified by DNA sequencing. HEK293 cells were purchased from the National Collection of Authenticated Cell Cultures (Shanghai Institutes for Biological Sciences, China), and cultured in Dulbecco’s Modified Eagle Medium (Gibco, USA) supplemented with 10% fetal bovine serum (FBS) (Gibco, USA), 1% penicillin-streptomycin, and 1% GlutaMAX™ (Gibco, USA). Cells were cultured at 37°C in a humidified atmosphere of 5% CO_2_ and 95% air. Plasmids were transfected into cells using a calcium phosphate transfection reagent. Unless otherwise stated, all other compounds were purchased from Sigma-Aldrich (USA).

Recordings of hP2X1, hP2X2, and hP2X3 receptor currents were performed using a conventional whole-cell patch configuration, as previously described (62). For conventional whole-cell recordings, the pipette solution comprised (in mM) 120 KCl, 30 NaCl, 0.5 CaCl_2_, 1 MgCl_2_, 10 HEPES, and 5 EGTA (pH 7.4, adjusted with Tris-base). Specifically, hP2X3 currents were recorded via perforated patch-clamp technique using nystatin to prevent current rundown. The nystatin (0.15 mg/mL) (Sangon Biotech, China) perforated intracellular solution contained (in mM) 55 KCl, 5 MgSO_4_, 75 K_2_SO_4_, and 10 HEPES (pH 7.4, adjusted with Tris-base). HEK293 cells were recorded after 36 hours of transfection using an Axopatch 200B amplifier (Molecular Devices, USA) with a holding potential of -60 mV at room temperature (25 ± 2°C). Current data were sampled at 10 kHz, filtered at 2 kHz, and analyzed by pCLAMP 10 (Molecular Devices, USA). HEK293 cells were bathed in standard solution (SS) containing (in mM) 2 CaCl_2_, 1 MgCl_2_, 150 NaCl, 5 KCl,10 HEPES, and 10 glucose (pH 7.4, adjusted with Tris-base). ATP and other drugs were dissolved in SS and perfused immediately onto the cell membrane during the recording period via Y-tubes.

### Molecular dynamics (MD) simulations

As previously described (62, 63), the energy-minimized models of hP2X3-sivopixant-ATP, hP2X3-sivopixant, hP2X3R-ATP, and apo hP2X3R were used as the initial structures for molecular simulations. A large 1-palmitoyl-2-oleoyl-sn-glycero-3-phosphocholine (POPC) bilayer (300 K), available in System Builder of Desmond (64), was built to generate a suitable membrane system based on the OPM database (https://opm.phar.umich.edu) (65), in which the TM domain of hP2X3 could be embedded properly. The hP2X3-sivopixant-ATP-POPC, hP2X3-sivopixant-POPC, hP2X3-ATP-POPC, and apo hP2X3-POPC systems were dissolved in simple point charge (SPC) water molecules. Counter ions were then added to compensate for the net negative charge of the system. NaCl (150 mM) was added into the simulation box that represents background salt at physiological condition. The DESMOND default relaxation protocol was applied to each system prior to the simulation run. Briefly, (1) 100 ps simulations in the NVT (constant number (N), volume (V), and temperature (T) ensemble with Brownian kinetics using a temperature of 10 K with solute heavy atoms constrained; (2) simulations were performed for 12 ps in the NVT ensemble using a Berendsen thermostat at 10 K with small time steps and solute heavy atoms constrained. (3) 12 ps simulations in the NPT (constant number (N), pressure (P), and temperature (T) ensemble using a Berendsen thermostat and barostat for 12 ps simulations at 10 K and 1 atm, with solute heavy atoms constrained; (4) 12 ps simulations using a Berendsen thermostat and Barostat at 300 K and 1 atm, with solute heavy atoms constrained; (5) 24 ps simulations using a Berendsen thermostat and Barostat at 300 K and 1 atm, with no constraints. After equilibration, the MD simulations were carried out for about 0.5 µs. The long-range electrostatic interactions were calculated using the smooth particle grid Ewald method. The trajectory recording interval was set to 200 ps and the other default parameters of DESMOND were used in the conventional molecular dynamics (CMD) simulation runs (66). All simulations used the OPLS 2005 all-atomic force field (67–69), which is used for proteins, ions, lipids and SPC waters. The Simulation Interaction Diagram (SID) module in DESMOND (62, 70) was used to explore the interaction analysis between SIV/ATP and hP2X3. All MD simulations were performed in DELL T7920 armed by NVIDIA TESLTA K40C or CAOWEI 4028GR armed by NVIDIA TESLTA K80. The simulation system was prepared, trajectory analyzed and visualized on a CORE DELL T7500 graphics workstation with 12 CPUs.

## Data availability

The atomic coordinates of the human P2X3 structures in the Protein Data Bank under accession codes 21FG (ATP- and sivopixant-bound, closed) [http://doi.org/10.2210/pdb21FG/pdb] and 21DX (ATP-bound, desensitized) [http://doi.org/10.2210/pdb21DX/pdb], respectively. Cryo-EM maps were deposited in the Electron Microscopy Data Bank (EMDB) under accession codes EMD-67624 (ATP- and sivopixant-bound, closed) [https://www.ebi.ac.uk/pdbe/entry/emdb/EMD-67624] and EMD-67603 (ATP-bound, desensitized) [https://www.ebi.ac.uk/pdbe/entry/emdb/EMD-67603], respectively. All other relevant data are included in the paper or its supporting information, or are deposited in Mendeley Data (https://doi.org/10.17632/crwytdnsdt.1).

## Author contributions

C.G. and M.H. designed research; Z.Z., D.W., X.Z., Y.G., C.S. and J.C. performed research; Z.Z., D.W., X.Z., Y.G., H.X., X.T., J.Z., Y.Y., C.G. and M.H. analyzed data; and Z.Z., D.W., C.G. and M.H. wrote the paper.

## Acknowledgments

We thank the staff scientists at the Cryo-EM Facility of the School of Life Sciences, Fudan University, for technical assistance with cryo-EM data collection. This work was supported by funding from the National Natural Science Foundation of China to M.H. (32471247, 32271244, and 32411540020) and C.G. (82474171). This work was also supported by the funding of the Postdoctoral Fellowship Program of CPSF to D.W. (GZC20252599) and the Open Research Fund of State Key Laboratory of Genetics and Development of Complex Phenotypes (No. SKLGDP2502, M. H.).

