## Supplementary material for "Cryo-EM reveals the structural basis of subtype-specific, noncompetitive inhibition of the human P2X3 receptor": Table S1 and Figure S1-6

### Supplementary Figure Legends

**Figure S1. Cryo-EM analysis.**

(A) Representative cryo-EM image of human P2X3 particles. (B) Representative 2D class averages. (C-G) For the sivopixant- and ATP-bound human P2X3 receptor: (C) Gold-standard Fourier shell correlation (FSC) curves for resolution estimation. (D) Angular distribution of the particles used for the final map. (E-G) Side view (E), top view from the extracellular side (F), and bottom view from the cytoplasmic side (G) of the cryo-EM density map colored according to the local resolution, estimated using CryoSPARC. (H-L) For the ATP-bound human P2X3 receptor: (H) Gold-standard FSC curves for resolution estimation. (I) Angular distribution of the particles used for the final map. (J-L) Side view (J), top view from the extracellular side (K), and bottom view from the cytoplasmic side (L) of the cryo-EM density map colored according to the local resolution, estimated using CryoSPARC.

**Figure S2. Cryo-EM data processing workflow.**

All the processing steps were performed using CryoSPARC.

**Figure S3. Comparison between cryo-EM and predicted structures.**

(A, B) Superimposition of the previously-predicted DDTPA-bound P2X3 structure (A) and the Boltz-2-predicted sivopixant-bound P2X3 structure (B) onto the cryo-EM structure of sivopixant- and ATP-bound P2X3 viewed from the extracellular side. The cryo-EM structure is shown in surface representation (upper right panel) or in ribbon representation (lower right panel) together with the superposed predicted structures (gray). The confidence score, protein ipTM, ligand ipTM, and complex pLDDT predicted by Boltz-2 were 0.81, 0.84, 0.53, and 0.80, respectively.

**Figure S4. AlphaFold3 prediction.**

(A, B) The predicted heterotrimer structure formed by one P2X2 subunit and two P2X3 subunits (AlphaFold3, ipTM=0.77) superposed onto the P2X3 structure shown in gray. In A, the gray chain superposed onto the yellow chain is the P2X2 subunit, while the gray chain superposed onto the blue chain is the P2X3 subunit. In B, the gray chain superposed onto the yellow chain is the P2X3 subunit, while the gray chain superposed onto the blue chain is the P2X2 subunit.

**Figure S5. Overall structural stability during the MD simulations.**

(A-H) MD simulations using the sivopixant- and ATP-bound structure with both retained (A, E, F), ATP deleted (B, G), sivopixant deleted (C, H) and both deleted (D) as starting models. The plots of the root mean square deviations (RMSDs) for Cα atoms (A-D) and the RMSD values of the atoms in sivopxant (E, G) and ATP (F, H).

**Figure S6. Allosteric pocket of P2X receptors**

(A-D) Close-up view of the ligand binding sites of the sivopixant- and ATP-bound P2X3 structure (this study) (A), of the camlipixant-bound P2X3 structure (PDB ID: 9BPC) (B), of the BAY-1797-bound P2X4 structure (PDB ID: 9BQI) (C), and of the JNJ-54175446-bound P2X7 structure (PDB ID: 8Z0Z) (D).

**Table S1. Cryo-EM data collection, refinement and validation statistics**

| | P2X3 with Sivopixant and ATP (EMDB:EMD-67624)(PDB: 21FG) | P2X3 with ATP (EMDB:EMD-67603) (PDB: 21DX) |
|---|---|---|
| **Data collection and processing** | | |
| Magnification | 130000× | 130000× |
| Voltage (kV) | 300 | 300 |
| Electron exposure (e–/Å$^2$) | 40 | 40 |
| Defocus range (μm) | -1.2 to -1.8 | -1.2 to -1.8 |
| Pixel size (Å) | 0.959 | 0.959 |
| Symmetry imposed | C3 | C3 |
| Initial particle images (no.) | 8747713 | 8747713 |
| Final particle images (no.) | 46550 | 71855 |
| Map resolution (Å) | 3.34 | 2.95 |
| FSC threshold | 0.143 | 0.143 |
| Map resolution range (Å) | 2.07 to 26.15 | 2.09 to 33.14 |
| **Refinement** | | |
| Initial model used (PDB code) | 5SVJ | 5SVL |
| Model resolution (Å) | 3.34 | 2.95 |
| FSC threshold | 0.143 | 0.143 |
| Model resolution range (Å) | 2.07 to 26.15 | 2.092 to 33.142 |
| Map sharpening *B* factor (Å$^2$) | -83.6 | -85.0 |
| Model composition | | |
| Non-hydrogen atoms | 7521 | 7764 |
| Protein residues | 981 | 984 |
| Ligands | SIV: 3; ATP: 3; NAG: 9 | NAG: 9; ATP: 3 |
| *B* factors (Å$^2$) | | |
| Protein | 110.01 | 106.86 |
| Ligand | 115.99 | 101.18 |
| R.m.s. deviations | | |
| Bond lengths (Å) | 0.003 | 0.002 |
| Bond angles (°) | 0.582 | 0.512 |
| Validation | | |
| MolProbity score | 1.52 | 1.46 |
| Clashscore | 3.88 | 3.73 |
| Poor rotamers (%) | 0.00 | 0.74 |
| Ramachandran plot | | |
| Favored (%) | 95.08 | 95.71 |
| Allowed (%) | 4.92 | 4.29 |
| Disallowed (%) | 0 | 0 |

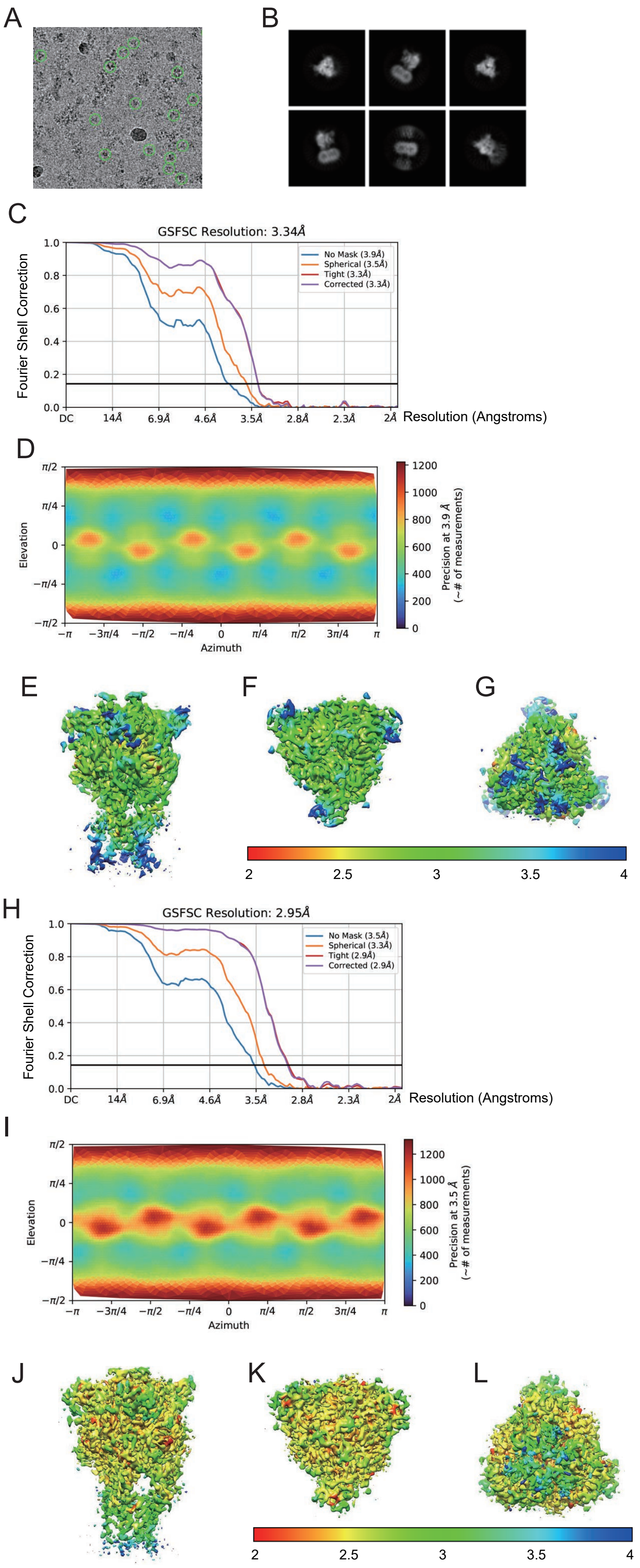
A
B
C
GSFSC Resolution: 3.34Å
Fourier Shell Correction
No Mask (3.9Å)
Spherical (3.5Å)
Tight (3.3Å)
Corrected (3.3Å)
DC
14Å
6.9Å
4.6Å
3.5Å
2.8Å
2.3Å
2Å
Resolution (Angstroms)
D
Elevation
Azimuth
Precision at 3.9 Å
(~# of measurements)
E
F
G
2
2.5
3
3.5
4
H
GSFSC Resolution: 2.95Å
Fourier Shell Correction
No Mask (3.5Å)
Spherical (3.3Å)
Tight (2.9Å)
Corrected (2.9Å)
Resolution (Angstroms)
I
Elevation
Azimuth
Precision at 3.5 Å
(~# of measurements)
J
K
L

Figure S1

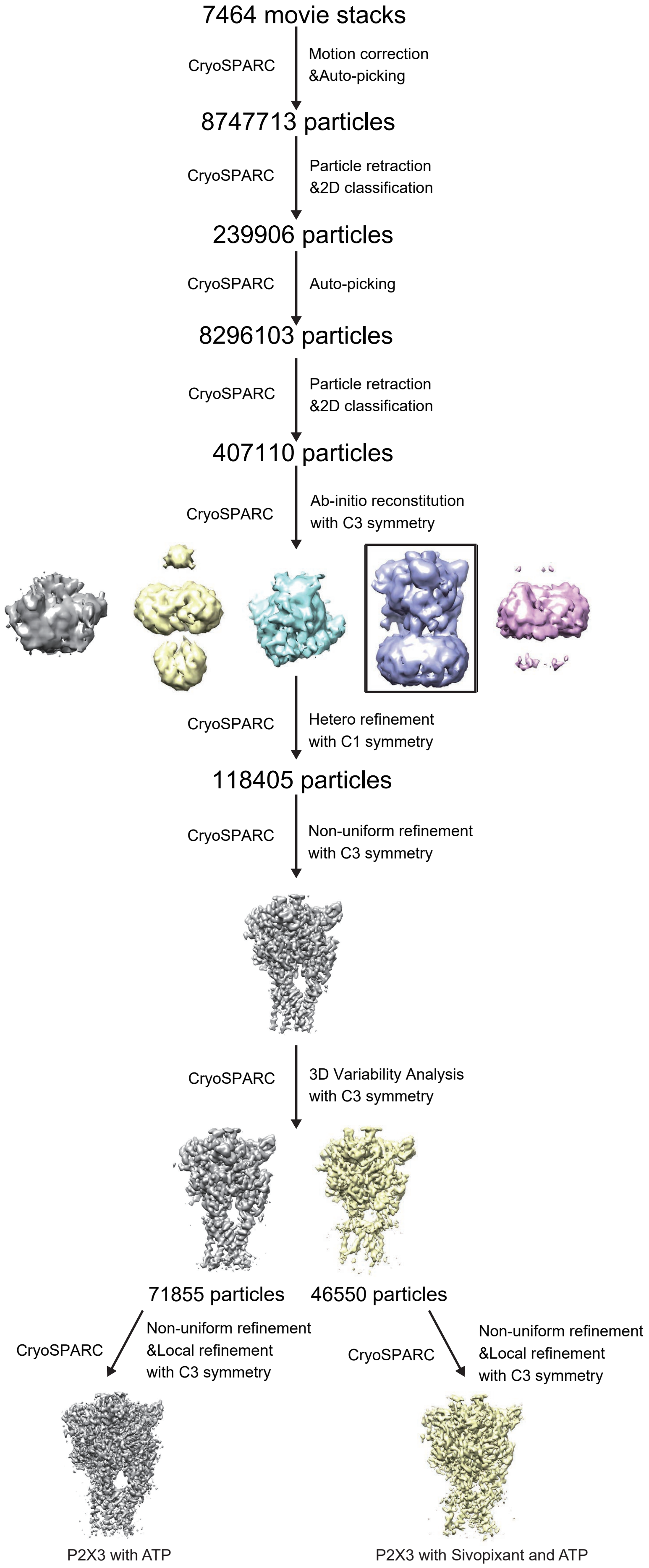
7464 movie stacks
CryoSPARC
Motion correction
&Auto-picking
8747713 particles
CryoSPARC
Particle retraction
&2D classification
239906 particles
CryoSPARC
Auto-picking
8296103 particles
CryoSPARC
Particle retraction
&2D classification
407110 particles
CryoSPARC
Ab-initio reconstitution
with C3 symmetry
CryoSPARC
Hetero refinement
with C1 symmetry
118405 particles
CryoSPARC
Non-uniform refinement
with C3 symmetry
CryoSPARC
3D Variability Analysis
with C3 symmetry
71855 particles
46550 particles
CryoSPARC
Non-uniform refinement
&Local refinement
with C3 symmetry
CryoSPARC
Non-uniform refinement
&Local refinement
with C3 symmetry
P2X3 with ATP
P2X3 with Sivopixant and ATP

Figure S2

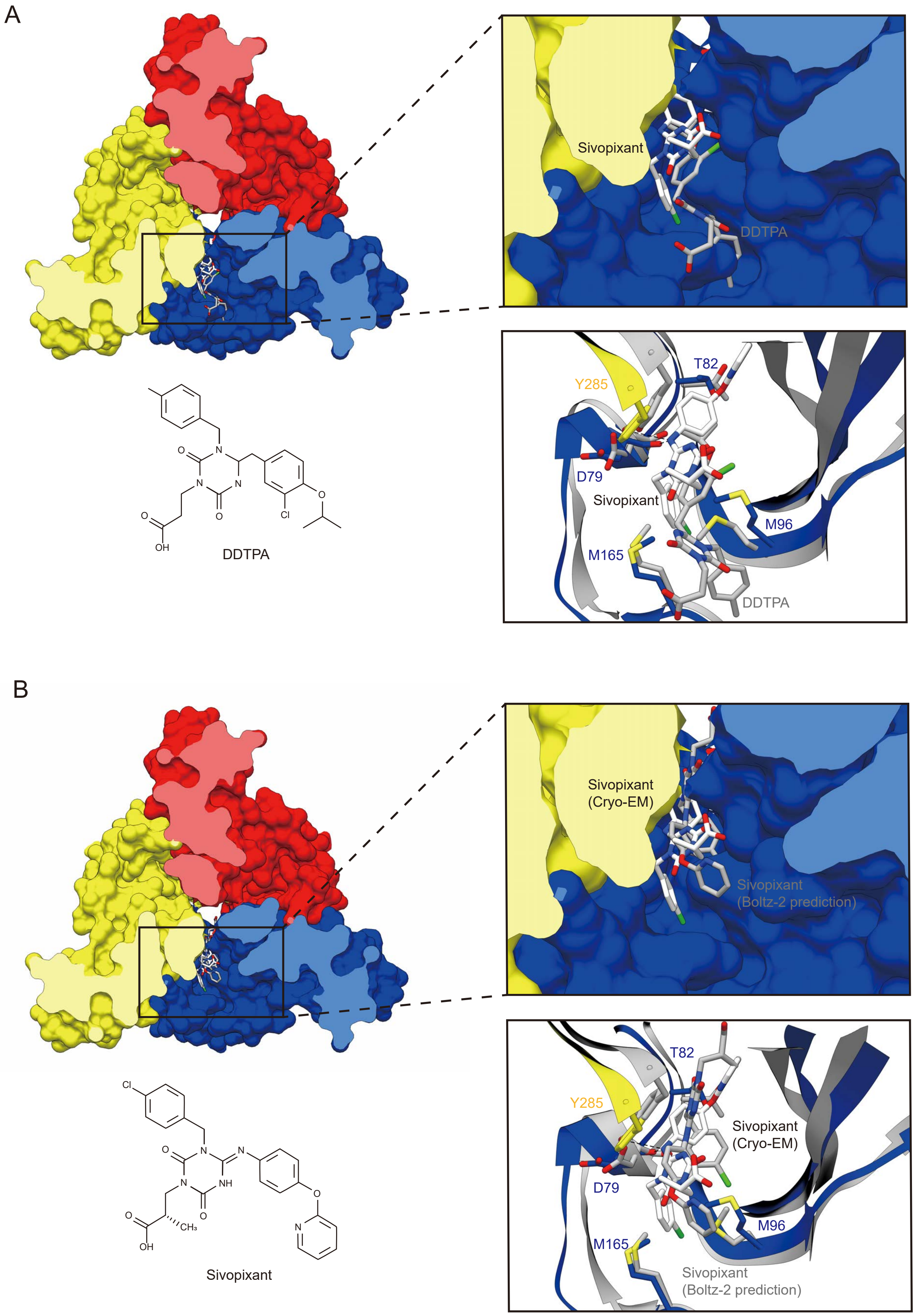
A
Sivopixant
DDTPA
T82
Y285
D79
Sivopixant
M96
M165
DDTPA
DDTPA
B
Sivopixant
(Cryo-EM)
Sivopixant
(Boltz-2 prediction)
T82
Y285
Sivopixant
(Cryo-EM)
D79
M96
M165
Sivopixant
(Boltz-2 prediction)
Sivopixant

Fiigure S3

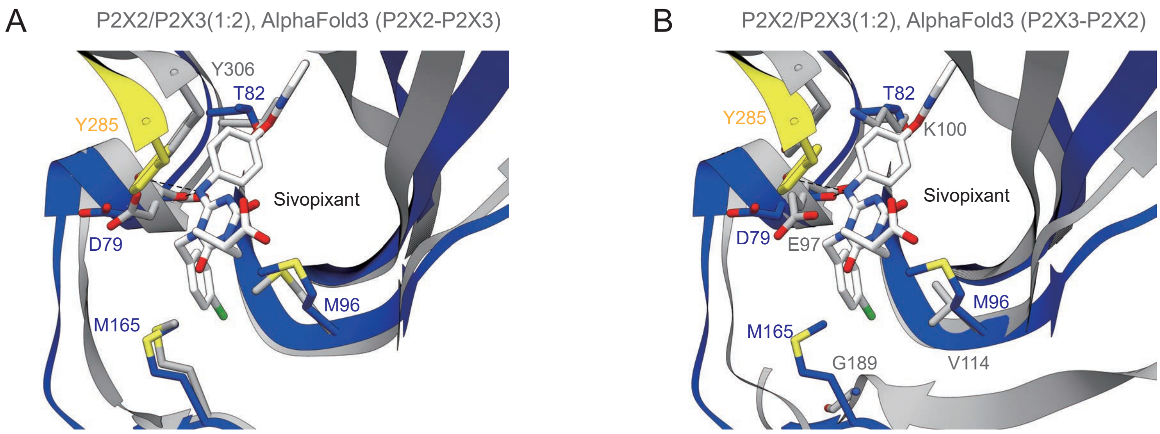

A
P2X2/P2X3(1:2), AlphaFold3 (P2X2-P2X3)
Y306
T82
Y285
Sivopixant
D79
M96
M165
B
P2X2/P2X3(1:2), AlphaFold3 (P2X3-P2X2)
T82
Y285
K100
Sivopixant
D79
E97
M96
M165
G189
V114


Figure S4

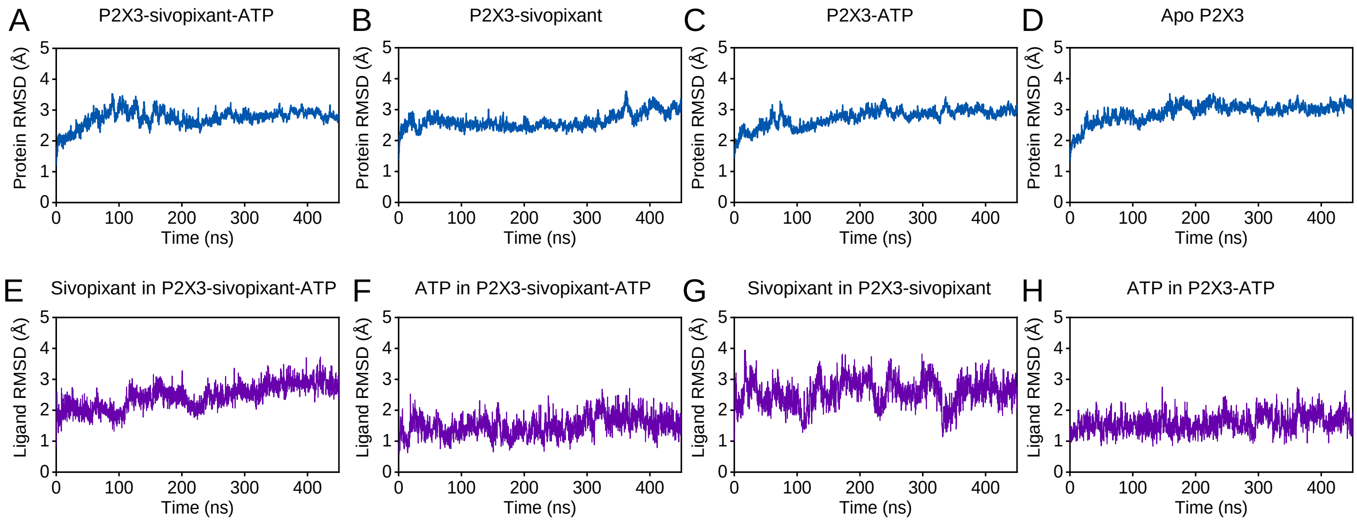
A
P2X3-sivopixant-ATP
B
P2X3-sivopixant
C
P2X3-ATP
D
Apo P2X3
Protein RMSD (Å)
Time (ns)
E
Sivopixant in P2X3-sivopixant-ATP
F
ATP in P2X3-sivopixant-ATP
G
Sivopixant in P2X3-sivopixant
H
ATP in P2X3-ATP
Ligand RMSD (Å)
Time (ns)

Figure S5

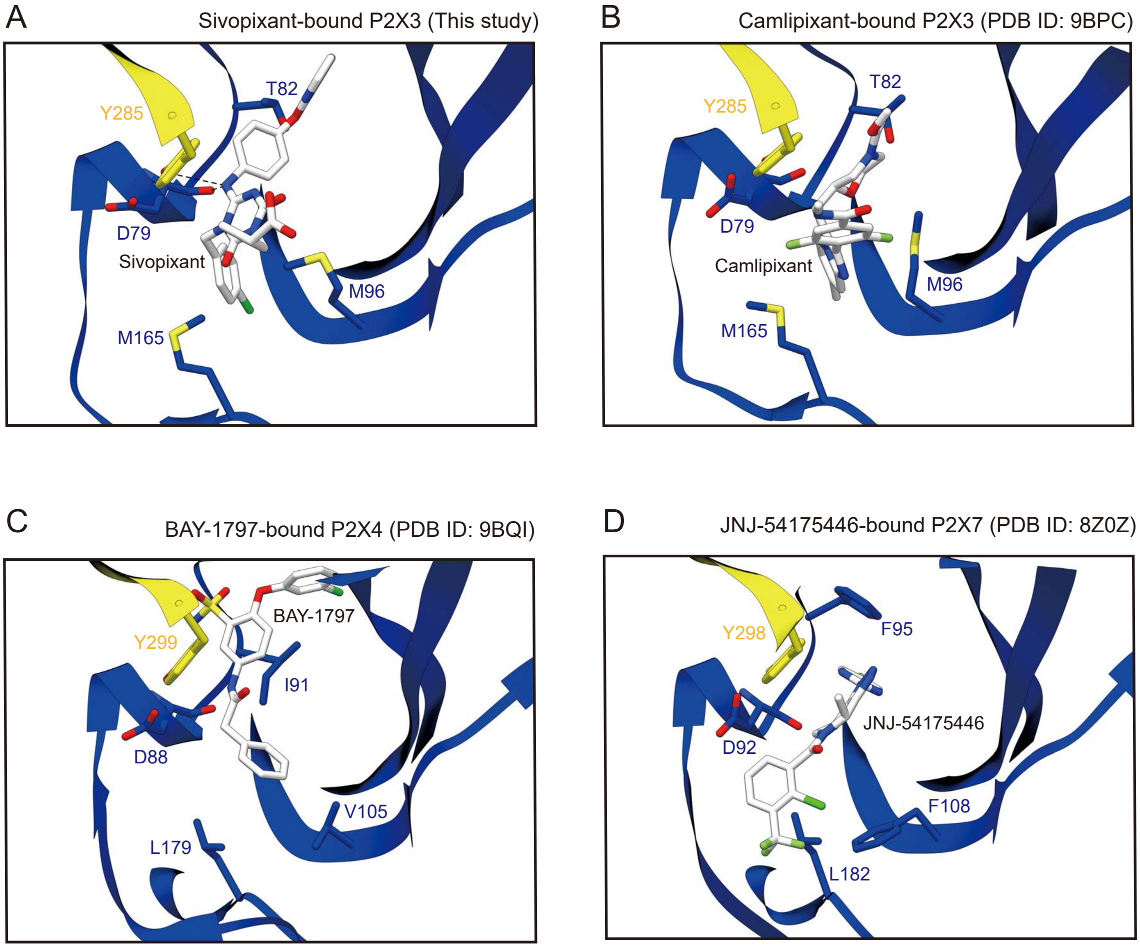
A
Sivopixant-bound P2X3 (This study)
Y285
T82
D79
Sivopixant
M96
M165
B
Camlipixant-bound P2X3 (PDB ID: 9BPC)
T82
Y285
D79
Camlipixant
M96
M165
C
BAY-1797-bound P2X4 (PDB ID: 9BQI)
BAY-1797
Y299
I91
D88
V105
L179
D
JNJ-54175446-bound P2X7 (PDB ID: 8Z0Z)
Y298
F95
JNJ-54175446
D92
F108
L182

Figure S6
